# Analyzing how habitat degradation drives extinction dynamics using physiologically-structured population models

**DOI:** 10.64898/2026.05.13.649732

**Authors:** Kenichi W. Okamoto, Virakbott Ong, Sergio A. Balaguera-Reina, Dat Dinh Phuc

## Abstract

Elucidating how habitat degradation facilitates extinction is critical for effective conservation efforts. Here, we propose integrating physiologically-structured population models into stochastic population viability analyses to assess how differing consequences of habitat degradation interact to drive extinction dynamics in a focal population. Using the isolated spectacled caiman *Caiman crocodilus* population/ecomorph from the Apaporis River as a case study, we find that threatening the resource base, which individuals increasingly rely upon, to outgrow vulnerable size ranges and mature accelerates extinction. We also found that when habitat degradation impacts both the primary adult and juvenile resource bases, this can have marked synergistic effects on threatening population viability. By contrast, destroying nesting sites has only a small effect on accelerating the impact of deteriorating prey availability. Through integrating community-level feedback between habitat degradation/change and population dynamics/structure, our approach provides a comparative framework for assessing the relative importance of distinct mechanisms through which habitat degradation ultimately drives extinction risk.

## Introduction

Few processes have as detrimental an effect on biodiversity as reduced quality and quantity of an organisms’ habitats (Sih et al. 2000; Carrete et al. 2009; Butchart et al. 2010; Turner 2010; Mantyka-pringle et al. 2012; Scanes 2018; Betts et al. 2022; Zhao et al. 2024). Among other mechanisms, habitat fragmentation caused by anthropogenic land conversion (Newburn et al. 2005; Boyd et al. 2015; Watson et al. 2016; Martin et al. 2020; Lindenmayer et al. 2023), habitat deterioration through accelerated extraction of keystone species (Estes et al. 2016; Wolf and Ripple 2016; Ripple et al. 2016; Ceballos et al. 2020; Woodward et al. 2021), and the introduction of pollutants accompanying infrastructure expansion, natural resource use, and urban and peri-urbanization (Peterson and Schulte 2016; Simkin et al. 2022) can all upend vital ecosystem services and expedite biodiversity loss (Young et al. 2016; Chase et al. 2020; Cafaro et al. 2022; Dasgupta and Levin 2023).

Several studies have assessed the impacts of habitat deterioration, fragmentation, and destruction on aggregate measures of biodiversity (e.g., total species richness, species compositions among functional or taxonomic groupings, Chao et al. 2019; Magurran et al. 2019; Chao et al. 2021; Tekwa et al. 2023). However, biodiversity loss ultimately results from the local extirpation of individual populations. Thus, characterizing how anthropogenic changes in species habitats impact extinction risks for particular populations presents a critical question for conservation biologists. Demography in particular is key to linking reduced habitat suitability and availability, on the one hand, to local extirpation, on the other. This is because we can leverage our considerable understanding of how demography drives extinction risk (e.g., Lacy 1993; Lande 1988; Beissinger and Westphal 1998; Klausmeier 1998; Green 2003; Drake 2005; Wootton and Pfister 2013; Benson et al. 2016; Conde et al. 2019; Katzner et al. 2020) to explain how anthropogenic changes in processes operating at the ecosystem and community-level threaten population persistence.

Population viability analysis (PVAs) provides a widely adopted, powerful conservation tool to characterize how environmental changes, such as habitat destruction, cascade to demographic changes and thus ultimately drive extinction risk (e.g., Boyce 1992; Brook et al. 2000; Lacy 2019; Chaudhary and Oli 2020). Using basic demographic and life-history data (e.g., fecundity, aging, and mortality), PVAs characterize how population dynamics (and, ultimately, extinction risk) emerge from the birth and death of individuals. At heart, PVAs rely on process-based/mechanistic models (i.e., models based on the underlying biological first principles – e.g., Cuddington et al. 2013) to characterize how population sizes change stochastically from one time step to the next (e.g., White 2000; Beissinger and McCullough 2002). By running many iterations of these stochastic models, PVAs quantify the probability that a population persists after a given period, enabling conservation and population biologists to characterize the contributions of specific processes to a population’s persistence probability (e.g., Morris et al. 1999; Jaffré and Le Galliard 2016).

Nevertheless, a major challenge for using PVAs to assess habitat loss risk is integrating the downstream effects of (often nonlinear) interactions between individual organisms and their changing environment that underlie key shifts in demographic and life-history processes driving extinction risks (e.g., Chisholm and Wintle 2007; Sabo 2008; Rayfield et al. 2009; Wittmer et al. 2014; Gudmundson et al. 2015). Such interactions are particularly prominent when evaluating how anthropogenic habitat changes affect life-history processes. Development through different life stages presents a case in point. Individuals do not develop identically (e.g., Lomnicki 1988; Nisbet et al. 2000; Bolnick et al. 2011; de Valpine et al. 2014). Rather, developmental variability emerges from a complex feedback loop involving resource availability, maturation schedules, physiological stress, etc… as well as inherent stochasticity (e.g., Violle et al. 2012; Moran et al. 2016; DeAngelis 2018; de Roos 2018). Although descriptive statistical models can sometimes account for these complexities (e.g., by regressing individual development rates on population size - e.g., Bjorndal et al. 2000; Bonenfant et al. 2009; Laver et al. 2012; Matthias et al. 2018; Zimmermann et al. 2018), by definition such models require data from past observations to construct. Such data can be difficult (at best) to collect, especially for rare taxa that are often the very subjects of PVAs (e.g., Simon 1969; Morris et al. 2002; Traill et al. 2007; Oppel et al. 2014; Chaudhary and Oli 2020). Moreover, if habitat loss and deterioration impose changes to local habitats beyond previously observed levels, the robustness of descriptive (or non-mechanistic) statistical models can be compromised, particularly given the nonlinear nature of the biotic and abiotic interactions involved. Hence, a framework linking habitat deterioration to extinction risk via demography will ideally allow key demographic and life-history processes to causally result from the (often) nonlinear interactions between individual organisms and the constituents of their environment.

To this end, we propose integrating stochastic implementations of physiologically-structured population models (PSPMs; Metz et al. 1986; de Roos 1997) into PVAs as a framework for causally evaluating extinction risks at the population level due to habitat degradation. Briefly, in PSPMs, an individual organism’s key demographic rates (reproduction and mortality) result from their physiological state (e.g., body size, energy reserves; Metz et al. 1986). Population dynamics, in turn, emerge from the aggregate effects of birth and death events across individuals. Thus, PSPMs characterize how changes in population states (density, size distribution, etc…) result from nonlinear feedbacks between environmental factors and individual condition (e.g., de Roos and Persson 2001). We therefore suggest that PSPMs can provide a mechanistic framework linking habitat condition to population viability. We illustrate the potential for using stochastic implementations of PSPMs in population viability analysis through a case study to evaluate how accelerating habitat degradation and destruction affect the risk of extinction of the unique spectacled caiman *Caiman crocodilus* population inhabiting the Apaporis River in Colombia (e.g., Medem 1955; Balaguera-Reina 2019; Balaguera-Reina et al. 2020; listed under the US Endangered Species Act as *C. crocodilus apaporiensis*). Using our proposed approach to PVAs, we analyze how varying drivers of habitat degradation differentially impact individuals in a population, thereby mechanistically linking anthropogenic habitat disruptions to demographic and population change and, ultimately, population extinction risk.

## Materials and Methods

### The physiologically-structured population model

We consider a single, size-structured, sexually- and seasonally-reproducing heterotrophic population with overlapping generations. We model the population’s biotic and abiotic factors in the environment to also potentially exhibit non-seasonal and secular dynamics. The population’s size and composition change as individuals in the population grow, reproduce, and die. Individual survivorship, somatic growth, and reproduction, in turn, all depend on individual physiological states. These physiological states are modeled to ultimately result from individuals in our population interacting with multiple, potentially dynamic resources in their habitat. Thus, by modeling the environment’s change over time, habitat degradation can be characterized as reductions in the availability of resources (in the very broadest sense) upon which individuals in the population depend on to survive and reproduce. This allows us to link anthropogenic perturbations to the population’s habitat to individual demographic outcomes, and thereby ultimately to population viability.

We begin by describing how we model the physiological states of individuals in the population. Following, e.g., Persson et al. (1998), somatic mass among individuals in the population is partitioned into two compartments: reversible mass, which consists of mass (e.g., gonads, lipid tissue) that can be starved away, and irreversible mass, which consists of mass that cannot be starved away (e.g., bones, certain organs). Individuals consume resources, which are metabolized and become energetic reserves, to be converted into somatic mass. Energetic reserves, in turn, are allocated into the two components of somatic mass according to individual condition - i.e., the relative availability of reversible mass. As in Persson et al. (1998), we model irreversible mass to not increase until an individual has sufficient reversible mass available. Once a condition threshold *q_·_* is exceeded, newly consumed energetic reserves can be allocated towards growth in both reversible and irreversible mass. As adults that have undergone maturation can allocate resources towards both storage and reproductive tissues, the condition threshold itself potentially differs for juveniles and adults. These considerations suggest the following series of equations describing changes in individual *j*’s physiological state:

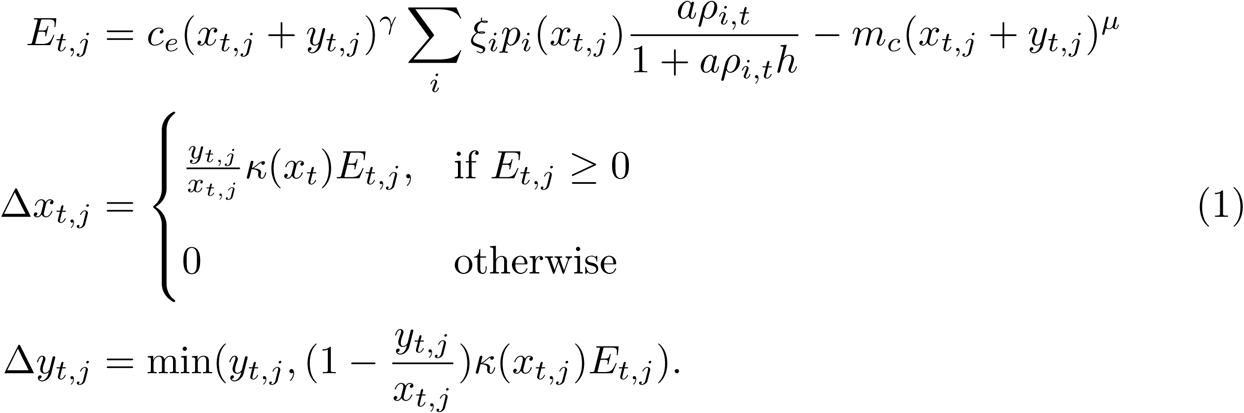

In eqs. (1), the individual’s state variables are *E_t,j_*, which represents the mass consumed (i.e., energetic gain) by an individual at time *t* and their irreversible mass *x_t,j_* and *y_t,j_*. *c_e_* describes the conversion efficiency of consumed biomass and *γ* is an allometric exponent characterizing how total food consumption scales according to total mass *x_t,j_* + *y_t,j_*. Individuals of total mass *x_t,j_*+ *y_t,j_* are further assumed to pay an allometric metabolic cost with allometric scalar *m_c_* and allometric exponent *µ*. The amount of food an individual can consume depends, in turn, on the availability *ρ_i,t_* of resource category *i* in the habitat at time *t*. The consumption rate for resource *i* at availability *ρ_i,t_* is driven by a maximum rate *a* and a quantity *h* that determines the resource availability at which consumption is halved. To enable common units in terms of mass available for somatic growth, we characterize resource availability *ρ_i,t_*in units of maximum *B_i_* energetic biomass of resource *i* available to the focal consumer population. Thus, *aρ_i,t_/*(1 + *aρ_i,t_h*) ∈ [0, 1) is a fraction controlling how the availability of resource *i* relative to some maximum *B_i_* impacts consumed biomass. As a consequence, the parameters *a, h* in eqs. (1) have somewhat different interpretations than the attack rate and handling time, respectively, commonly used in Holling Type-II functional responses (e.g., Holling 1959). The function *p_i_* potentially allows individuals of different sizes to consume different resources i.e., undergo an ontogenetic niche shift (e.g., Werner and Gilliam 1984; Nakazawa 2015). In practice, for many species the shape of the ontogenetic niche shift is largely determined by components of structural mass (e.g., gape size) instead of reversible mass (e.g., Claessen and Dieckmann 2002; De Roos and Persson 2013). In the presence of an ontogenetic niche shift, the logistic function *p_i_*(*x_t,j_*) = 1*/*(1 + *e^αi^*^+*β*^*^ixt,j^*) provides a convenient representation of the fraction of the diet of an individual with irreversible mass *x_t_* consisting of resource *i*; *α* governs the proportion of the diet consisting of resource *i* for very small individuals, and *β* determines the size at which half the diet consists of resource *i*. If more than two resource groups are available, then *p_i_*(*x_t,j_*) can be further scaled to represent the fraction of each resource group consumed at a given size.

Once consumed mass is determined across all resource categories, the available energy is allocated to irreversible or reversible mass based upon the function *κ* = 1*/*(*q_·_*+ *q*^2^) that differs between mature adults, who must allocate reserves to reproduction as well as storage, and juveniles, who do not. We follow, e.g., Persson et al. (1998) in assuming that the size *s_m,·_* at maturation, which can vary for males and females, depends on developmental processes associated with growth in irreversible mass (e.g., body length). Finally, if resources are rare, *E_t_* can potentially become negative as individuals incur a metabolic cost. If *E_t_*is negative, then irreversible mass cannot increase, and any available reversible mass diminishes to cover the metabolic cost.

The dynamics of individual physiological state described in model (1) drive population growth through their impacts on mortality and reproduction. An individual’s reproductive potential *r* is determined by *y_t_*−*x_t_q_A_*, which represents the surplus mass (above the adult-specific condition threshold *q_A_*) available for reproductive activities. In females, this surplus mass is converted to new individuals according to *rp_e_/*(*q_J_ ɛ* + *ɛ*), where *ɛ* represents the irreversible mass of eggs (for oviparous species) or newborns (for viviparous species) and *p_e_* is the fraction of viable eggs or neonates. Although the surplus mass available to adult females determines the number of neonates, we note that males can also potentially expend surplus mass for reproduction (e.g., to defend mating territories, provide parental care, etc…).

The other way in which the dynamics of model (1) governs population dynamics is through their effects on mortality. We model three sources of mortality: (i) size-specific mortality, (ii) starvation-induced mortality and (iii) background mortality. Because in many populations individuals with smaller irreversible mass tend to have lower survivorship (e.g., due to predation suffered at vulnerable size classes; Deevey 1947; Claessen et al. 2004), we model the size-specific mortality risk using the logistic expression 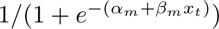, where *α_m_*determines the background mortality risk of very small individuals and *β_m_*the marginal change in mortality risk as individuals grow. Starvation reduces survivorship exponentially according to 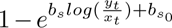, where the parameters *b_s_, b_s_*_0_ govern how rapidly deteriorating individual condition induces mortality. Finally, a baseline mortality *µ_s_* is imposed on all individuals to account for processes such as senescence. Individual mortality risk is thus the product of these differing sources of mortality.

### Habitat degradation effects on population viability

The physiologically-structured population model presented in the previous section permits us to evaluate how different consequences of habitat destruction/ deterioration potentially threaten population viability. We focus on two major classes of resource degradation frequently cited in the literature: reduced prey available for consumption (e.g., Ryall and Fahrig 2006; Tylianakis et al. 2007; Jellyman et al. 2014; Martinson and Fagan 2014) and reductions in suitable breeding or nesting sites (e.g., Dolman and Sutherland 1995; Hallworth et al. 2021).

Reduced prey resources available for consumption can result from the proximate consequences of habitat destruction and deterioration (e.g., Wolf and Ripple 2016). For instance, sewage runoffs, siltation, agricultural pollutants, and overharvesting of biological resources can all disrupt the basal food web communities upon which species occupying higher trophic positions depend (e.g., Deegan, L. A. and Buchsbaum, R. 2005; Burdon et al. 2020). Moreover, if prey exhibit limited mobility compared to predators (e.g., Malmqvist 2002; Gehring and Swihart 2003; Hillaert et al. 2020), over relevant spatial scales their abundances may prove more sensitive to, e.g., habitat fragmentation (e.g., Bascompte and Solé 1998; Macdonald et al. 1999; Swihart et al. 2001). Prey are also typically smaller than their predators (e.g., Cohen et al. 1993; Brose et al. 2006), which can lead to prey populations being more vulnerable than predator populations to habitat degradation (e.g., Ripple et al. 2017).

The second mechanism for how habitat degradation threatens population viability that we explore is the reduction in nesting or breeding sites (e.g., Mainwaring et al. 2017). For many animals, breeding or nesting sites can often be identified, leading to ready measurements of changes in their availability. Conservation concerns associated with the destruction of sites used for reproduction have been widely documented, including insects (e.g., Flockhart et al. 2017; Wagner 2020), molluscs (e.g., Rothschild et al. 1994; Wilberg et al. 2011; Arkhipkin et al. 2015), amphibians (e.g., Cushman 2006), mammals (e.g., Harwood 2001; Johnstone et al. 2014), crustaceans (e.g., Lejeusne and Chevaldonné 2006; Smith et al. 2017), non-avian reptiles (e.g., Marchand and Litvaitis 2004; McClenachan et al. 2006; Böhm et al. 2016), fish (e.g., Deegan, L. A. and Buchsbaum, R. 2005; Einum et al. 2008; Munday et al. 2008; Dahlke et al. 2018), and birds (e.g., Dolman and Sutherland 1995; Stephens et al. 2004).

Thus, the combination of potentially density-independent and density-dependent consequences of habitat degradation presents considerable challenges to conservation efforts. Because they can characterize how somatic growth depends on resource availability, physiologically-structured population models facilitate integrating both density-independent and density-dependent mechanisms into our understanding of population dynamics and, hence, extinction risks. However, the effects of habitat degradation on density-dependent processes are governed, in part, on how recruitment of new individuals into the population changes over time. Thus, because of the central role of breeding and nest site availability plays in the recruitment process, we examine how the direct loss of such sites impacts extinction risk, both alone and when coupled with changes to prey resource availability.

As a case study, we illustrate how these two consequences of habitat degradation can be incorporated into population viability analyses using the unique spectacled caiman population that inhabits the Apaporis River. This distinctive ecomorph Balaguera-Reina et al. 2022 is currently included in Appendix I of the Convention on International Trade in Endangered Species of Wild Fauna and Flora (CITES; CITES Secretariat 2014) to reduce the impact of trade on this sensitive population. The habitat of this ecomorph is restricted to the middle Apaporis River basin in Colombia, South America (Medem 1955; Balaguera-Reina 2019). An analysis of Mitochondrial DNA (mtDNA) markers in Balaguera-Reina et al. (2020) demonstrates the population of the putative *C. c. apaporiensis* appears situated within the broader Amazonian *C. crocodilus crocodilus* lineage of the *C. crocodilus /C. crocodilus yacare/C. yacare* species complex (Roberto et al. 2020), at least as mtDNA is concerned. Nevertheless, the *C. crocodilus* population in the middle Apaporis River basin is of notable conservation interest due to its distinctive ecomorphological characteristics and prey specificity (e.g., Medem 1955; US Fish and Wildlife Service 1976; Ayarzagüena 1984; Escobedo-Galván et al. 2015; Okamoto et al. 2015; Angulo-Bedoya et al. 2019; Balaguera-Reina et al. 2020; Falcón Espitia and Jerez 2021; Okamoto and Balaguera-Reina 2026).

While *C. crocodilus* as a whole may be adaptable to human-disturbed landscapes (e.g., Thorbjarnarson and Velasco 1999; Parks et al. 2024; but see Pereira et al. 2022), species distribution modeling in related *C. yacare*/*C. c. yacare* populations also suggests habitat degradation and other environmental changes associated with deforestation and climate change could meaningfully drive range contraction (Rodriguez-Cordero et al. 2022). Since the diminution of armed conflict in 2016, the Colombian Amazon has undergone dramatic and sustained deforestation (Clerici et al. 2020). Of particular concern from the perspective of habitat deterioration and destruction has been the rapid expansion of cattle ranching (Murillo-Sandoval et al. 2023) - for instance, Hoang et al. (2023) identified the geographic risk profile raising cattle presents to areas of conservation priority in Colombia as being among the highest in the world.

The consequences of habitat degradation described above can therefore potentially pose threats to the spectacled caiman population that inhabits the Apaporis River. For example, there is evidence the niche shift in this population may differ from other *C. crocodilus* populations Okamoto and Balaguera-Reina 2026. Hence, deteriorating prey availability can result from, among other drivers, increased human harvesting pressure on fish stocks and agricultural runoff associated with cattle ranching (e.g., Junk et al. 2007) - anthropogenic disturbances which, to be sure, can also affect prey availability beyond ichthyofauna. Furthermore, alterations to river morphology and flow, as well as land-use changes associated with deforestation have the potential to reduce nest site suitability in other crocodylians (e.g., Cavalier et al. 2022).

Here we integrate community ecology theory into our physiologically-structured population model (PSPM) to perform a population viability analysis for a localized unique population of spectacled caimans. In addition to its conservation interest, this population presents an attractive candidate case study for our proposed approach integrating PSPMs into PVAs. Like all crocodylian populations, the *C. crocodilus* population in the Apaporis River exhibits considerable size variation, with individuals growing from a little over 10 cm SVL shortly after hatching, to over 125 cm SVL as mature adults (Medem 1981). This size variation, in turn, has important implications for their community ecology (e.g., Ayarzagüena 1984; Okamoto and Balaguera-Reina 2026), and therefore linking physiologically and size-structured ecological processes to population persistence is particularly relevant in this system. Additionally, the biology of *C. crocodilus* and its close relatives has been the topic of fairly intense research for decades (e.g., Grigg and Kirshner 2015), in part due to their economic importance (e.g., Brazaitis et al. 1998; Thorbjarnarson and Velasco 1999). Consequently, plausible parameter estimates are available for constructing PSPMs for use in PVAs in this population. Fig. 1 summarizes the key processes the PSPM models, and how it links individual development to population and community dynamics. Our analyses focus on two major proposed mechanisms on how habitat degradation can threaten population viability: reduced prey availability and destruction in nesting sites.

**Figure 1.**
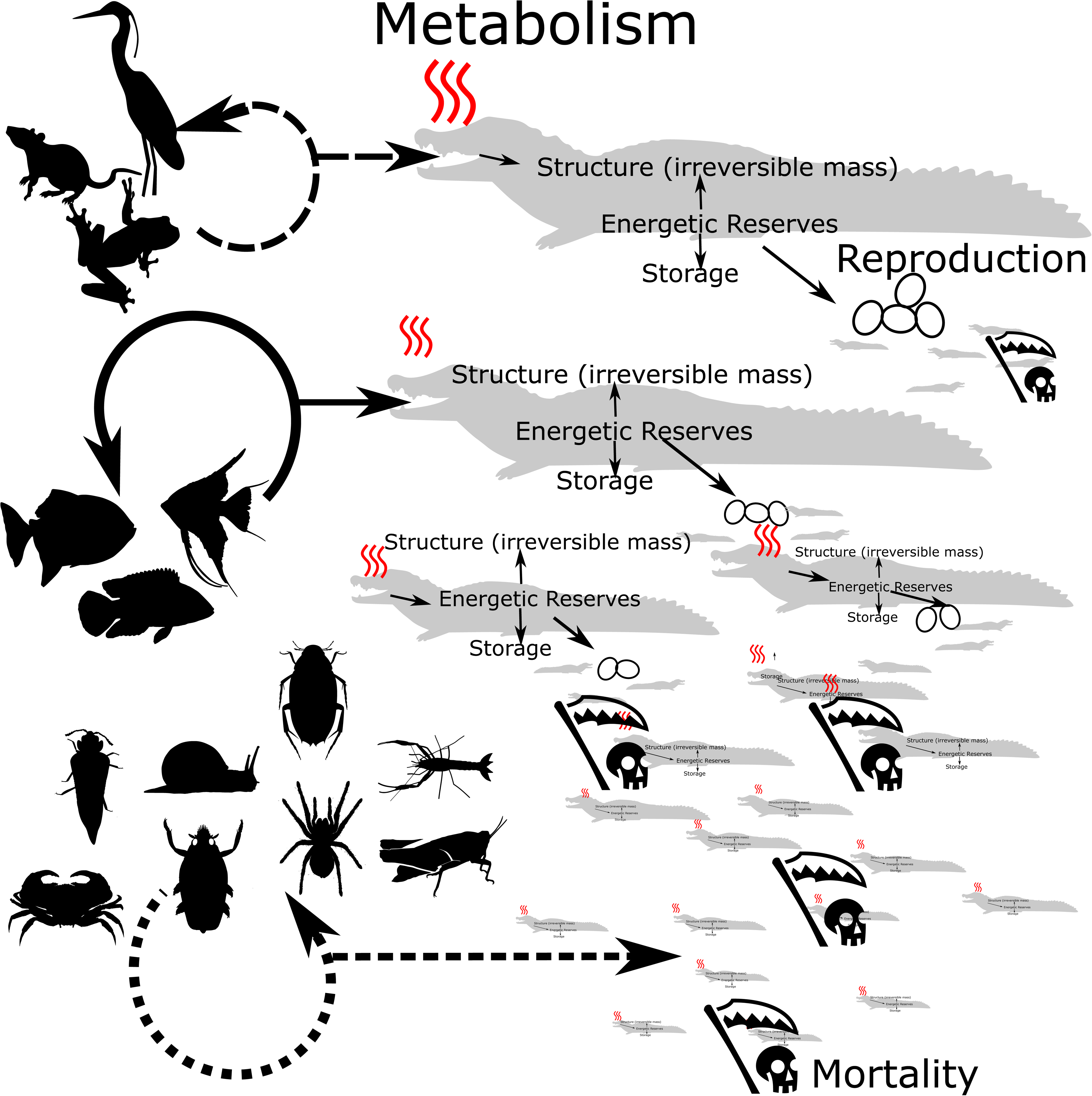
The key processes of the physiologically-structured population model and their relationships to population and community dynamics. Individuals consume prey, and the energetic surplus after metabolism is allocated towards irreversible (structural tissue) and reversible (storage and reproductive tissue) mass. Individual physiological states affect their life histories - i.e., somatic growth, mortality and reproductive schedules. Image credits: https://creativecommons.org/licenses/by-nc/3.0/ CC BY-NC 3.0: modified from Armin Reindl, FSock and JCGiron. CC0 1.0 Universal Public Domain Dedication: Hugo Gruson, Katy Lawler, Andy Wilson, Carlos Cano-Barbacil, Birgit Lang, T. Michael Keesey, Nathan J Baker, Martin R. Smith, Guillaume Dera and Stevan Traver.

To characterize the impact of habitat degradation on prey availability, we model the dynamics of resource *i* using Beverton-Holt recruitment dynamics (e.g., Beverton and Holt 1993). We adopt the Beverton-Holt recruitment model for two reasons. First, it can be derived under a wide range of underlying biological assumptions (e.g., Geritz and Kisdi 2004; Brännström and Sumpter 2005; de la Parra et al. 2013), and second, its dynamics in discrete time result from integrating the widely used logistic growth model in continuous time (Verhulst 1838). Thus, in the absence of habitat degradation, the dynamics of resource biomass (in units of consumable biomass for the focal population - here, the Apaporis River *C. crocodilus* population) are modeled as 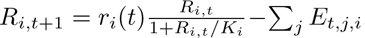, where *r_i_*(*t*) is the intrinsic growth rate in consumable biomass at time *t*, *K_i_* is a quantity characterizing density-dependence in resource biomass *i*’s growth and which affects the carrying capacity of consumable biomass of resource *i* in the habitat, and ∑*_j_ E_t,j,i_* the amount of resource *i*’s consumable biomass lost to consumption across all individuals *j* of the focal consumer population.

Combined with the PSPM presented earlier, this framework allows us to model the adverse effects of habitat degradation on food resources in two ways. First, we characterize the most proximate effect of habitat degradation as progressively reducing *K_i_* and thus the carrying capacity of prey group *i*’s consumable biomass. Second, deteriorating habitats have a notoriously destabilizing effect on population and community dynamics (e.g., Gonzalez et al. 2011). These destabilizing effects can result, for instance, from an increased likelihood of random catastrophes faced by populations (e.g., Lande 1993; Wilcox and Elderd 2003) but can also (at least temporarily) increase resource biomass, for instance, through the introduction of invasive species in disturbed habitats (e.g., Sakai et al. 2001; Koehn 2004; MacDougall and Turkington 2005; Schliserman et al. 2014; Chichorro et al. 2022). Thus, as a second complementary consequence to long-term reductions in resource availability, we model these effects of habitat degradation as destabilizing the intrinsic rate *r_i_*(*t*) of resource biomass growth by *r_i_* + *η_i_*(*t*)*r_i_*, where *η_i_*(*t*) ∼ *N* (0*, σ_i_*). We follow Agresti (2002) and Okamoto and Balaguera-Reina (2026) in partitioning *C. crocodilus* prey into three functional, taxonomic groups: invertebrates, fish, and tetrapods.

In contrast to prey resources, which are dynamic, we model nesting and breeding sites for the Apaporis River *C. crocodilus* population to be a constant resource over time in the absence of habitat degradation. When habitat destruction is present, available nesting and breeding sites decline. We model the scarcity of nesting sites to place strong constraints on the focal population’s recruitment as follows: when there are more nesting and breeding sites than there are reproducing females in the population, each such female is assumed to be able to secure a site. Otherwise, the probability that an individual reproducing female is able to obtain a breeding or nesting site is given by the number of nesting and breeding sites divided by the number of reproductive females. As habitat destruction and deterioration cause the availability of nesting sites to decline, population dynamics are impacted as more and more females are unable to reproduce.

Although reductions in nesting sites can reduce density-independent recruitment, simultaneously, they can potentially reduce density-dependent mortality resulting from, for instance, intraspecific resource competition for food. Nevertheless, as prey populations are also negatively impacted by habitat degradation, such releases from density-dependent constraints may prove transient. A key advantage of our approach is that we can readily analyze and quantify how these two potentially countervailing impacts of habitat degradation impact long-term population viability.

We examine how each of these three mechanisms interact to elevate extinction risks by varying the relative severity with which habitat destruction degrades the different resources. First, we consider how habitat degradation increases negative density-dependence and destabilizes the intrinsic growth rate separately for the prey categories primarily consumed by juveniles (i.e., invertebrates) and adults (fish and tetrapods) along a qualitative gradient of reduction in nest site availability from modest (10% reduction) to considerable (90% reduction). We thereby characterize how habitat degradation affecting prey availability for one life stage, but not another, interacts with nest site reduction to impact extinction risk. We then consider cases where habitat degradation causes a correlated destabilization of prey intrinsic growth and increased density-dependence across prey types at different life stages. Such correlated effects are likely if, for instance, one prey class (e.g., ichthyofauna) also consumes another prey class (e.g., aquatic and semiaquatic macro-invertebrates), in which case reduced population growth in the basal resource (e.g., invertebrates) can adversely affect both prey resource groups. We highlight how this strategy enables us to compare the effects on habitat degradation on density-independent and density-dependent processes not only for our focal population, but for its prey as well.

Table 1 summarizes the variables and parameters used in the model for our case study, and Table 2 summarizes the model equations discussed for our PSPM. Table 3 summarizes our characterization of the consequences of habitat degradation used in our analyses.

**Table 1:**
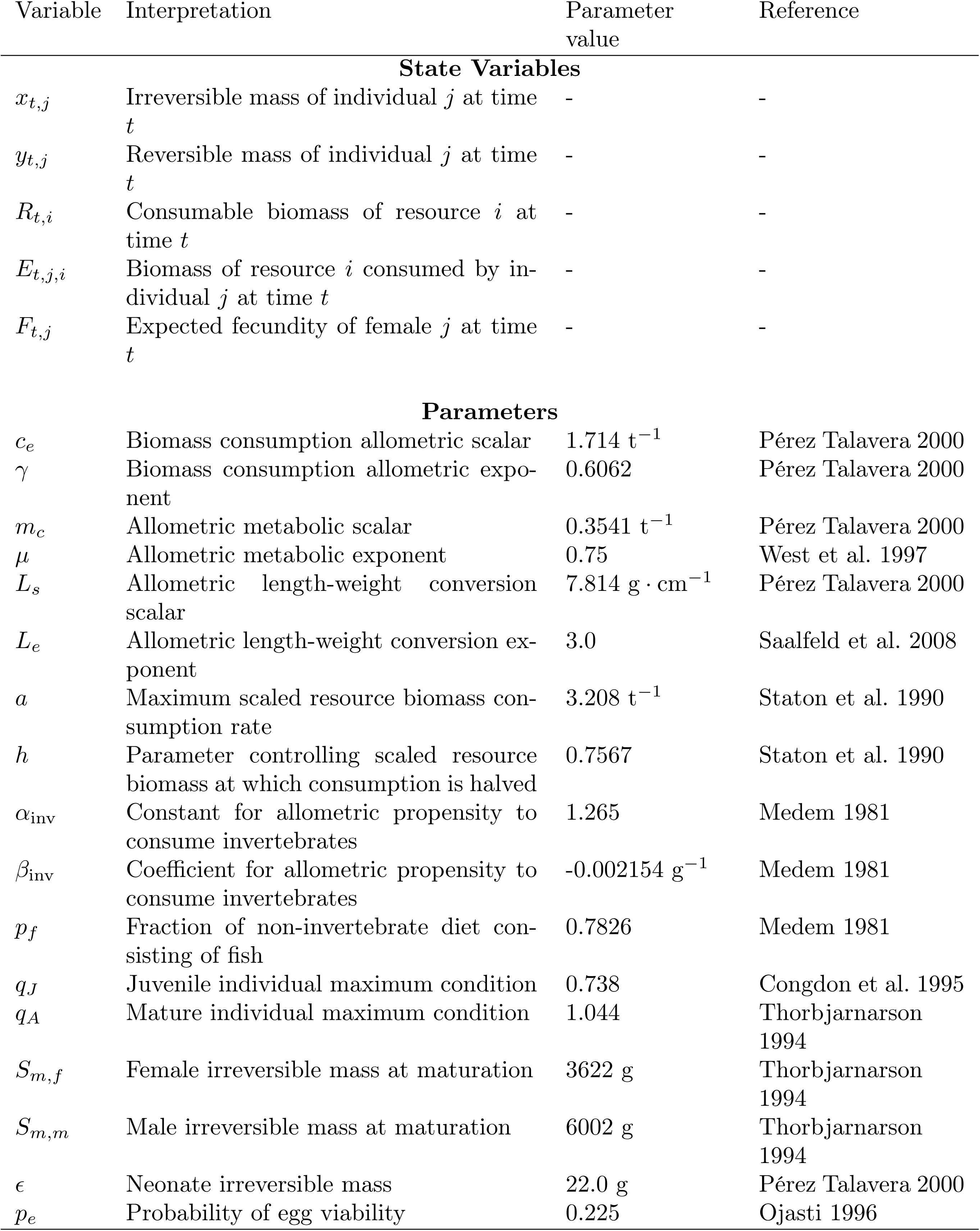
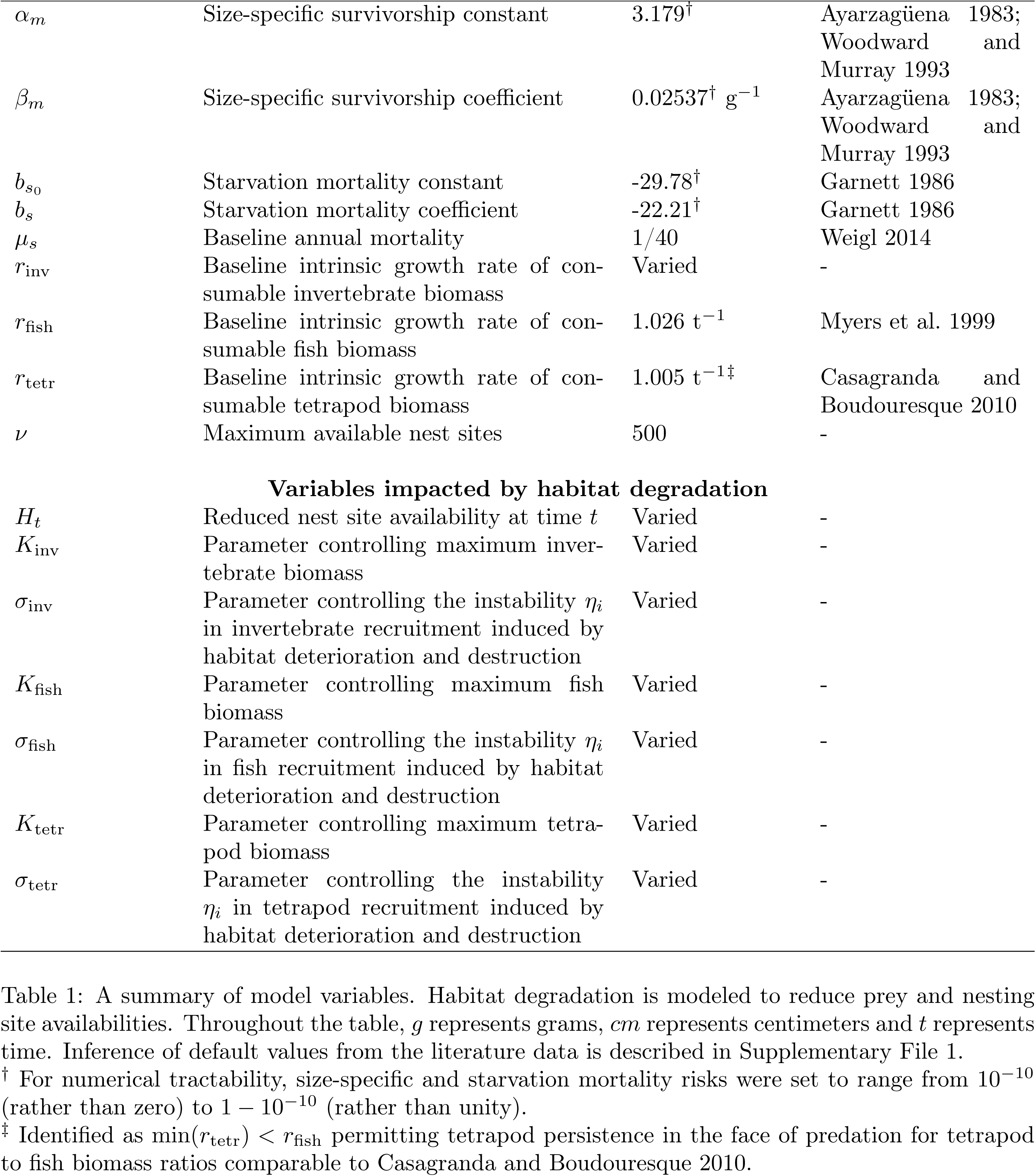
A summary of model variables. Habitat degradation is modeled to reduce prey and nesting site availabilities. Throughout the table, *g* represents grams, *cm* represents centimeters and *t* represents time. Inference of default values from the literature data is described in Supplementary File 1. † For numerical tractability, size-specific and starvation mortality risks were set to range from 10*^−^*^10^ (rather than zero) to 1 − 10*^−^*^10^ (rather than unity). ‡ Identified as min(*r*_tetr_) *< r*_fish_ permitting tetrapod persistence in the face of predation for tetrapod to fish biomass ratios comparable to Casagranda and Boudouresque 2010.

**Table 2:**
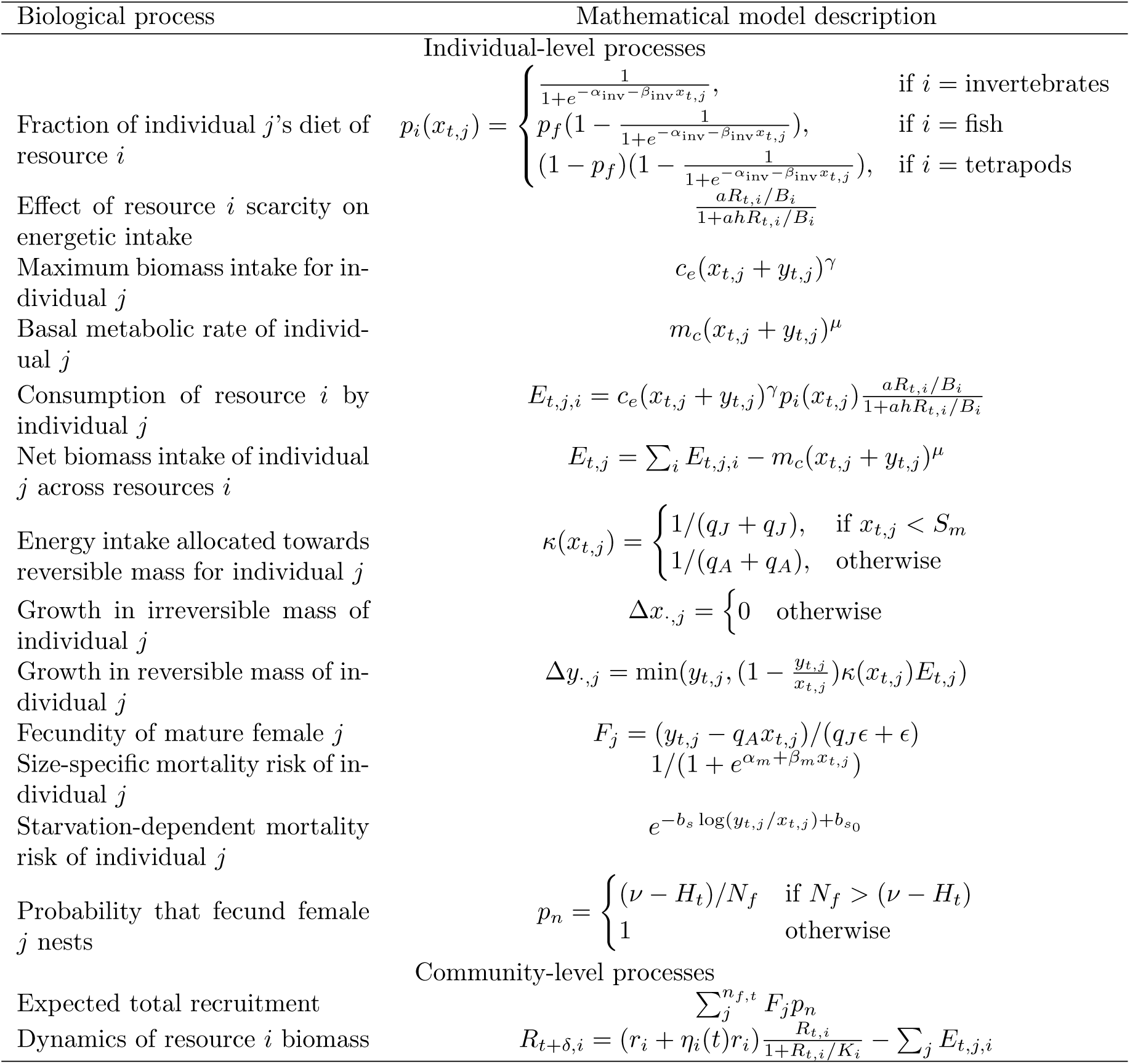
A summary of the processes modeled. The dynamics of the physiologically-structured population model (PSPM) emerge from the interaction between processes operating at the individual-level for all individuals and processes operating at the community level.

**Table 3:** Modeled mechanisms by which habitat degradation drives extinction risks. In the absence of any effects of habitat degradation (i.e., when all parameter values in this table are zero), the population does not go extinct. For nest site availability and reduced prey carrying capacities, habitat destruction is modeled to proceed linearly until the specified degradation level is reached at the end of the simulated time horizon.

| Table 3 |  |
| --- | --- |
| Effect of habitat degradation | Range |
| Nest site reduction as fraction of available nests | 0.1-0.9 |
| Reduction in invertebrate prey carrying capacity | 0-0.9999 |
| Reduction in vertebrate prey carrying capacity | 0-0.9 |
| Instability of invertebrate prey unconstrained biomass growth (as fraction of $r_i$) | 0-0.25 |
| Instability of vertebrate prey unconstrained biomass growth (as fraction of $r_i$) | 0-0.075 |

To perform population viability analyses, we implemented the PSPM as stochastic, individual-based realizations of the processes described in Tables 2-3 over a period of 1000 years following a 100 year burn-in period using the Simulating Phenotypic Evolution on General-purpose Graphics-processing units (sPEGG) framework (Okamoto and Amarasekare 2018). We analyzed extinction dynamics over this 1000-year period for 100 independent replicate runs for each scenario of habitat degradation in Table 3. Code underlying the analysis is publicly available at https://anonymous.4open.science/r/pspm_pva-8ECC and released under the GNU Public License v3 (Stallman 2007). The codebase includes a Jupyter/colab notebook (**?**) that requires minimal coding backgrounds which readers can readily run on their Google Drive.

## Results

Our key result is that the mechanisms by which habitat degradation drive extinction dynamics qualitatively differ depending on which prey type, or combination of prey types, are impacted by habitat loss.

For both prey groups (invertebrates consumed primarily by smaller individuals and vertebrates increasingly consumed as individuals grow), extinction accelerates sharply when habitat degradation strengthens the effects of density-dependence on prey recruitment (and hence reduces the prey carrying capacity; Figs. 2-5). Across scenarios, as habitat degradation reduces the prey group’s carrying capacity over about 85% of its original level, population extinction occurs rapidly within the first third to half of the simulated time horizon.

**Figure 2.**
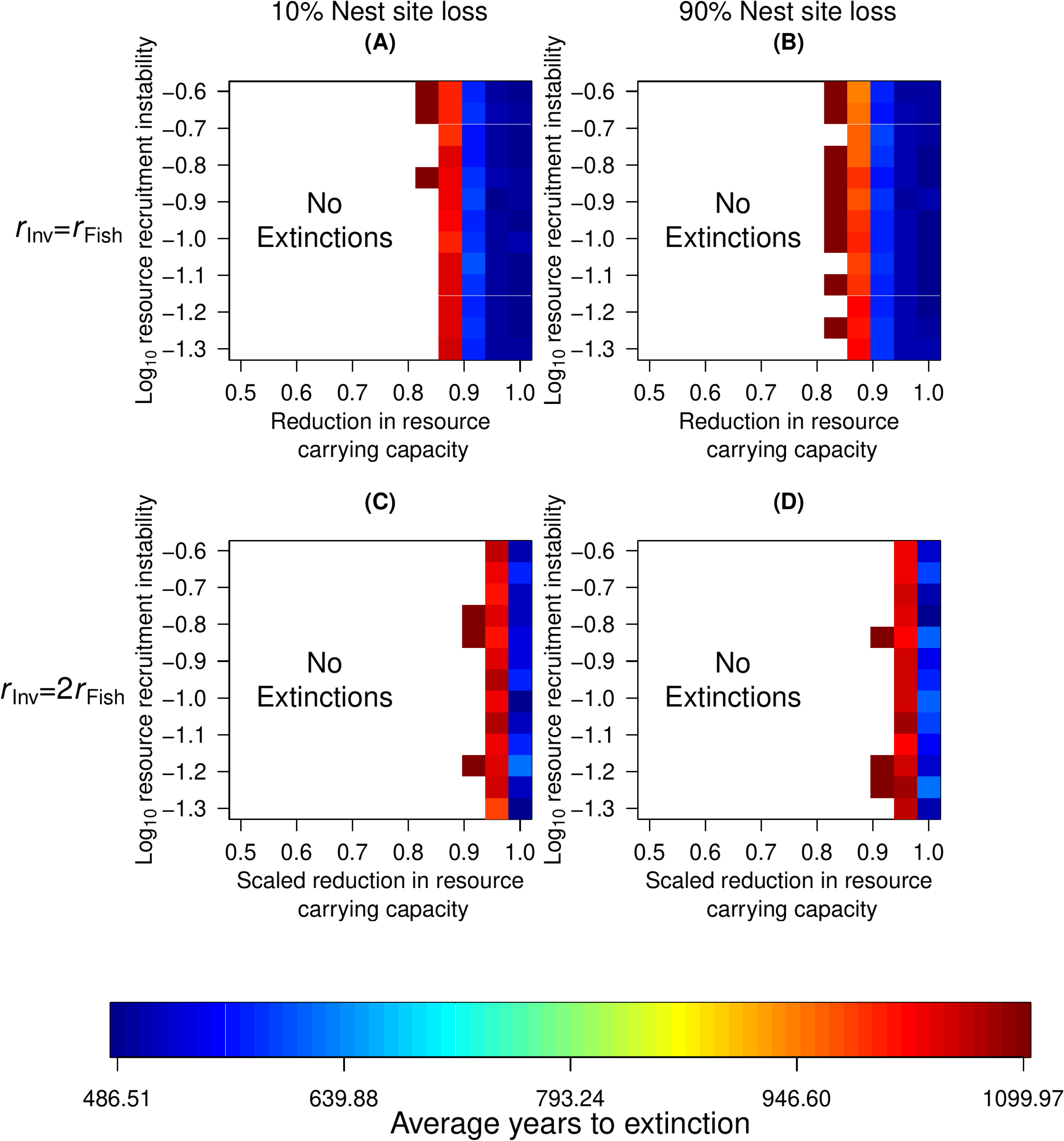
Average times to extinction (in simulated years) across 100 replicate simulations when habitat degradation impacts only consumable invertebrate biomass recruitment. Here, and in subsequent figures, *B_·_*= (*r_·_* − 1)*K_·_* and, at the start of the simulations and for the duration of the burn-in period, *K*_fish_ = *K*_tetr_ = 5 × 10^11^g and *K*_inv_ = 10*K*_fish_(e.g., Lindeman 1942); regions in white represent areas of parameter space where no extinctions occurred.

**Figure 3.**
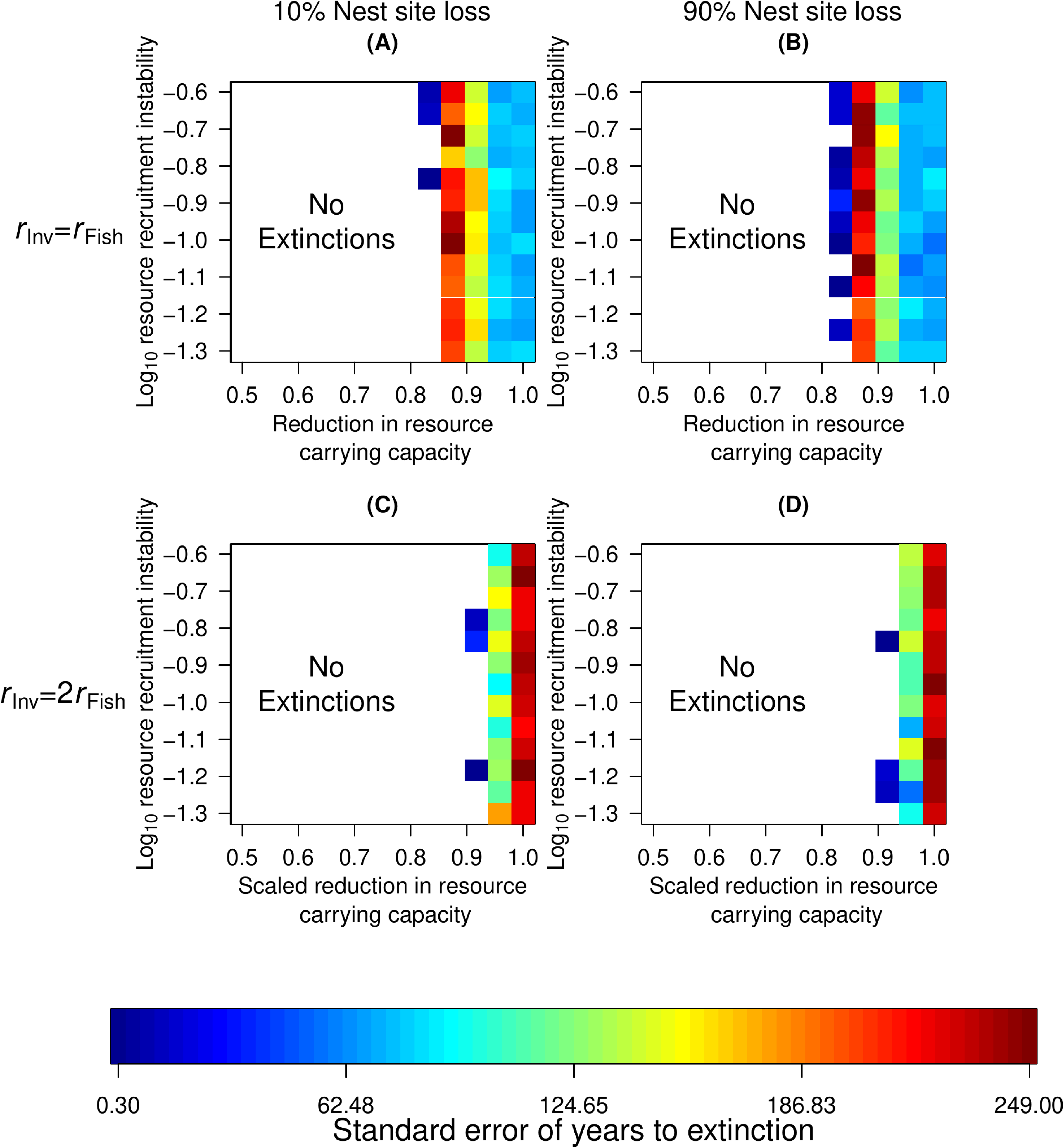
The standard error for the times to extinction (in simulated years) across 100 replicate simulations when habitat degradation impacts only consumable invertebrate biomass recruitment.

**Figure 4.**
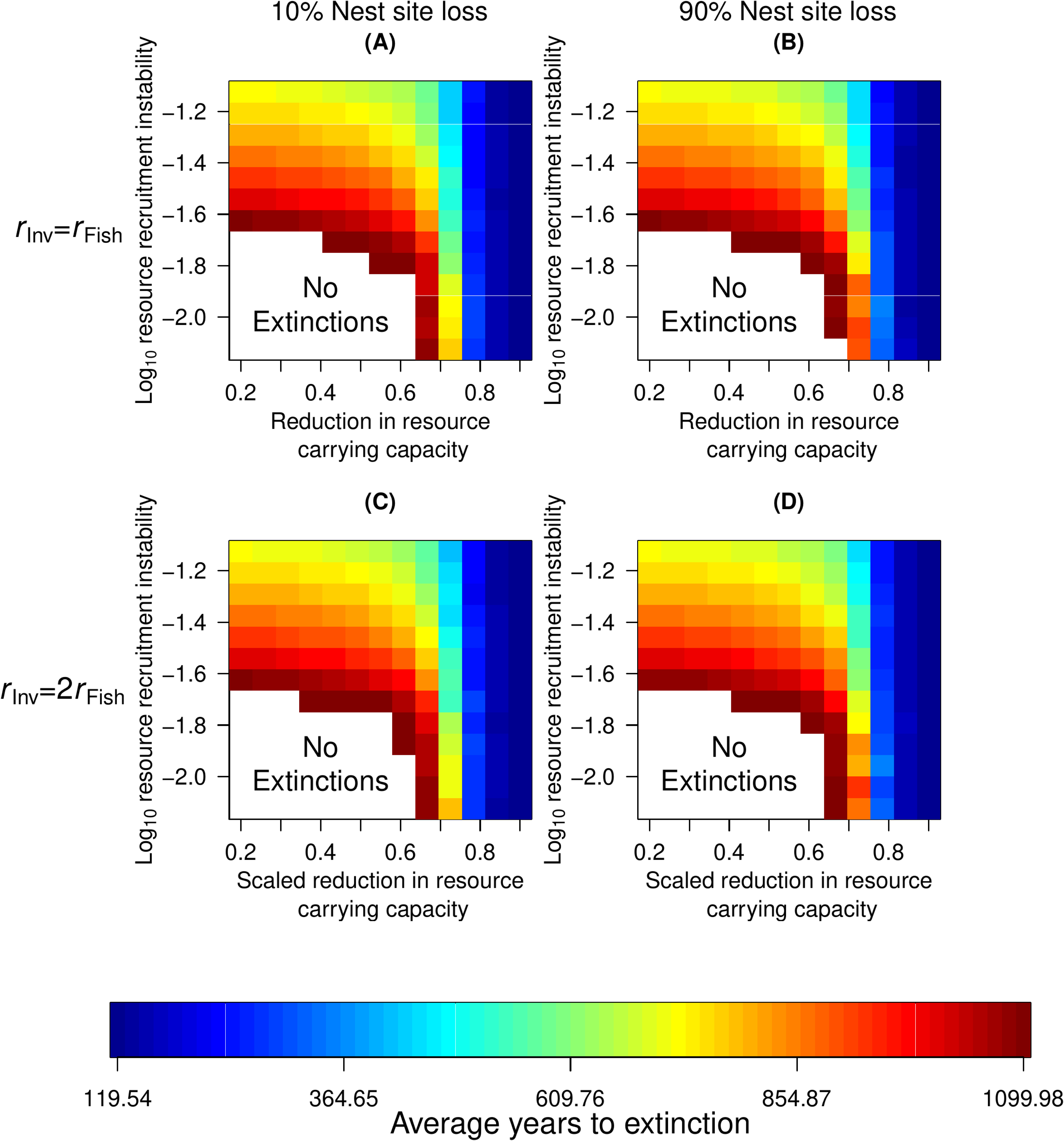
Average times to extinction (in simulated years) across 100 replicate simulations when habitat degradation impacts only consumable vertebrate (fish and tetrapod) biomass recruitment.

**Figure 5.**
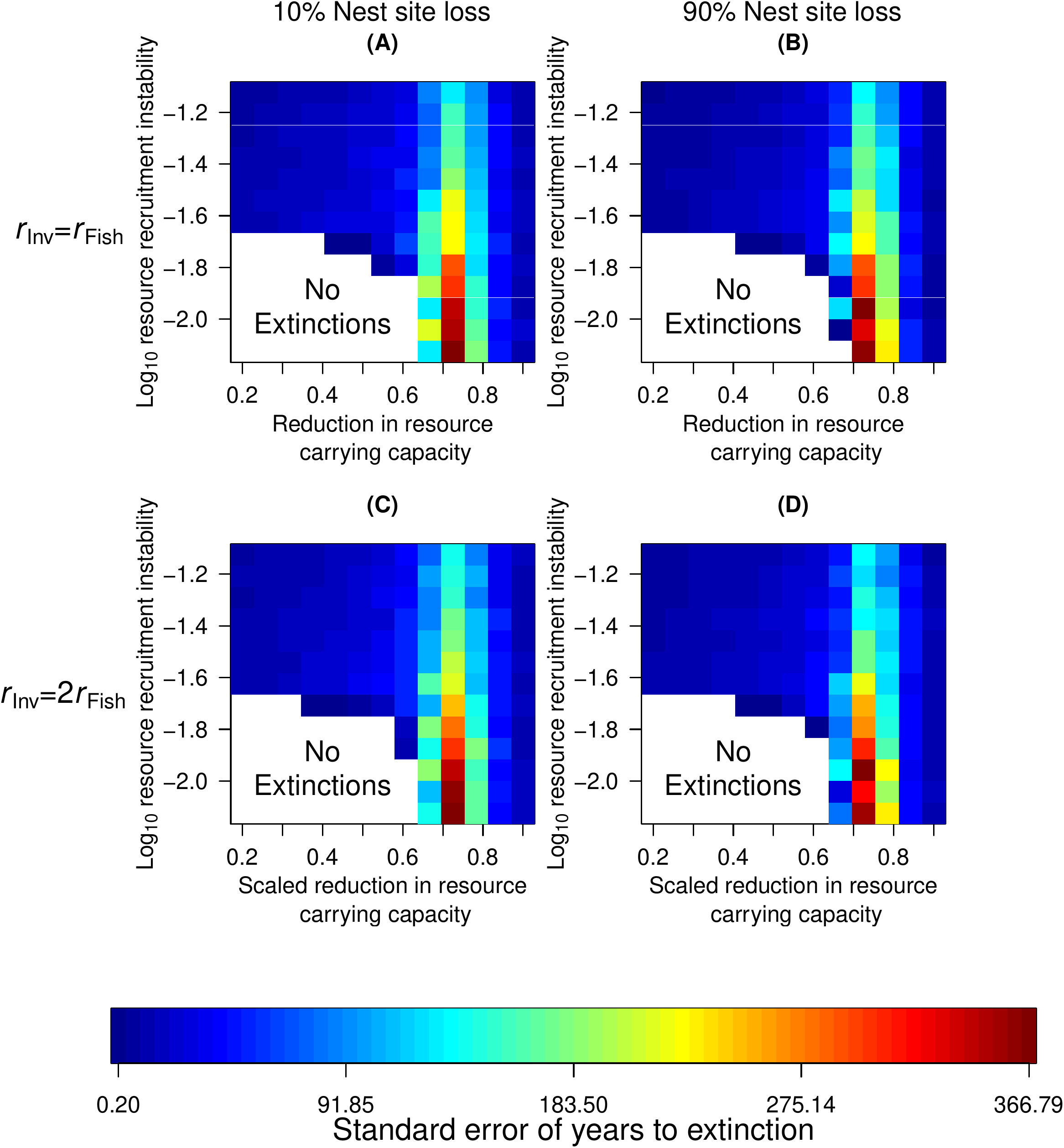
The standard error for the times to extinction (in simulated years) across 100 replicate simulations when habitat degradation impacts only consumable vertebrate biomass recruitment.

In contrast to reductions in the prey group’s carrying capacity, when habitat degradation impacts invertebrate, but not vertebrate, prey recruitment, destabilizing the prey group’s unconstrained growth rate does not have a strong effect on extinction dynamics over the range examined (Fig. 2). This result holds even when looking at higher order moments (Fig. 3). This is in contrast to the case when habitat degradation affects only vertebrate prey availability (Figs. 4-5). When vertebrate, but not invertebrate, prey recruitment is affected by habitat degradation and destruction, instability in the prey’s unconstrained growth mediates the effect of reductions in the carrying capacity on extinction risks (Figs. 4). In particular, as density-independent vertebrate prey recruitment is destabilized, the viability of the spectacled caiman population at the Apaporis River becomes more sensitive to reductions in the vertebrate prey’s carrying capacity. For instance, if habitat degradation causes the vertebrate prey’s unconstrained growth rate to be highly variable (around 0.08*r_·_*), even modest reductions in the vertebrate prey’s carrying capacity cause eventual extinction (Fig. 4, upper portion of all panels). Yet, when the vertebrate prey’s unconstrained growth rate is relatively stable in the face of habitat degradation (around 0.0075*r_·_*), even when the prey’s carrying capacity is substantially reduced, several replicate spectacled caiman populations persist for considerable durations (Fig. 5, bottom right corner of all panels).

Despite this contrast, destabilizing the invertebrate prey’s density-independent recruitment rate can still alter extinction dynamics when habitat degradation affects both prey groups. There, destabilized prey recruitment more readily causes extinctions at lower levels than when only vertebrate prey are impacted by habitat degradation (Fig. 4 vs. Fig. 6). Put differently, were destabilizing vertebrate density-independent recruitment alone driving extinction dynamics, we would have expected similar effects of destabilized unconstrained growth whether habitat degradation impacted invertebrate prey (Fig. 6) or not (Fig. 4).

**Figure 6.**
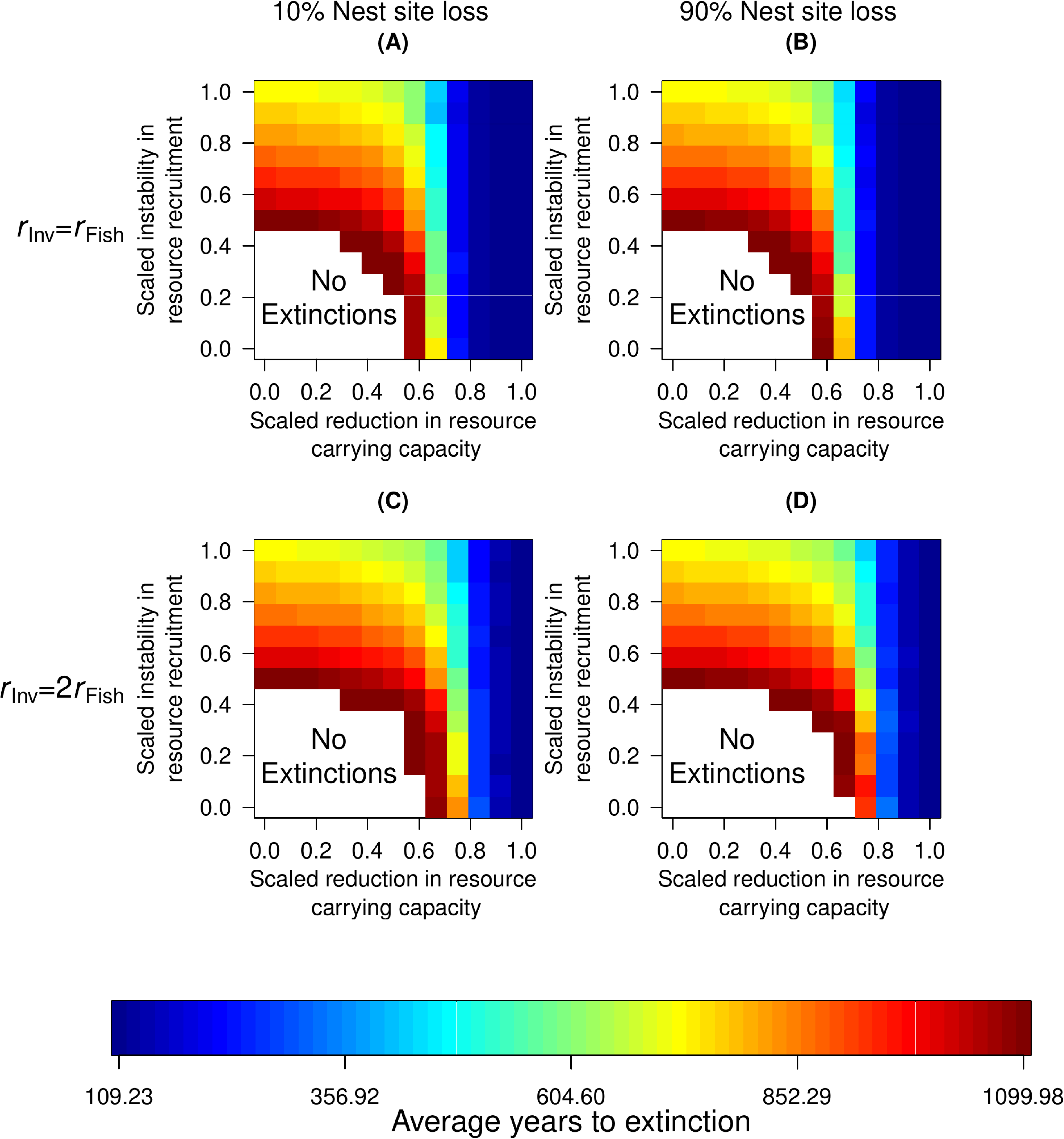
Average times to extinction (in simulated years) across 100 replicate simulations when habitat degradation impacts both prey group’s consumable biomass recruitment. Here, and in the subsequent figure, for each panel, the abscissa and ordinate are scaled to correspond to their respective ranges in Figs. 2-5 for both prey categories.

A further contrast to the cases where habitat degradation affects only one prey group is illustrated by the higher order moments of extinction dynamics (Fig. 7). There, we find consistently rapid extinctions when habitat degradation reduces both prey’s carrying capacity at the maximal levels analyzed. The only other scenarios where the time to extinction proved consistent at high levels of habitat degradation across replicate runs was when habitat degradation only impacted invertebrate prey and the unperturbed density-independent recruitment of invertebrate prey was comparable to that of fish (Fig. 2A-B).

**Figure 7.**
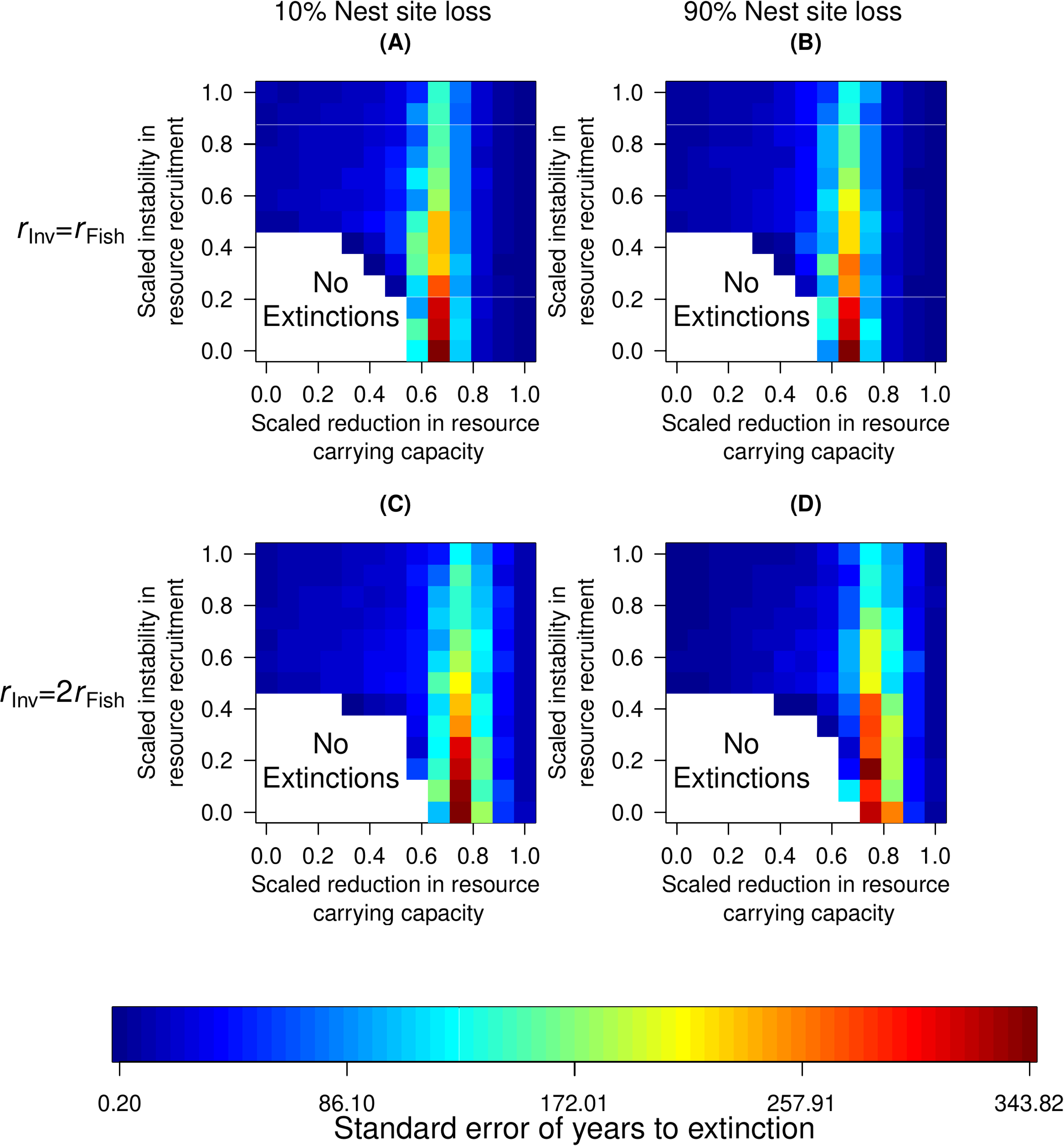
The standard error for the times to extinction (in simulated years) across 100 replicate simulations when habitat degradation impacts both prey groups’ consumable biomass recruitment.

Increasing the rate of nest site destruction consistently accelerated the time to extinction for given levels of reduced prey carrying capacity (Figs. 2-7, panels A and C vs. panels B and D). However, we note that the elevated loss of nesting sites proved to have a mainly quantitative, rather than qualitative, impact on extinction dynamics across scenarios for how habitat degradation affects different prey groups. Similarly, when the baseline density-independent recruitment rate of invertebrate biomass was larger than that of ichthyofauna, this enabled spectacled caiman populations from the Apaporis River to persist more readily under more severe conditions of habitat degradation (Figs. 2-7, panels A and B vs. panels C and D). Yet, as was the case for the destruction of nesting sites, the qualitative impacts of habitat degradation on extinction dynamics proved consistent across changes in density-independent invertebrate biomass growth rates.

## Discussion

Linking specific consequences of anthropogenic habitat changes to population dynamics is critical to understanding threats to biodiversity. This is because extinctions caused by habitat degradation are ultimately mediated by organismal traits and community-level feedbacks (Hanski 2011; González-Suárez et al. 2013; Chichorro et al. 2022). Here, we proposed and applied a mechanistic framework to elucidate how the interplay between the community-level effects of habitat degradation on biotic and abiotic resources can cascade through individual life histories to impact population viability. In this study, we proposed using physiologically-structured population models (PSPMs) to assess how distinct consequences of habitat degradation interact with each other to drive extinction dynamics.

Applying this approach to a unique and isolated population like the spectacled caiman found at the Apaporis River as a case study, we find that this population is especially vulnerable to habitat changes degrading the carrying capacity of the biotic resource base -vertebrate prey- upon, which individuals increasingly rely as they grow in size. When only the biotic resource individuals exploit at smaller sizes and earlier in life -invertebrate prey in our case study- is impacted by habitat degradation, considerable destruction in the resource’s carrying capacity must occur before extinction risks accelerate. We further find that the deleterious effects of reducing the juvenile resource’s carrying capacity appears generally invariant to the extent to which the prey group’s unconstrained recruitment rate is destabilized by habitat destruction.

Our results therefore present a meaningful advance on the existing literature by elucidating how reduced prey group availability at different size classes interacts with population heterogeneity to drive extinction risks. As spectacled caimans at the Apaporis River grow in size, they transition to exploiting vertebrate prey. Degradation in vertebrate prey availability can stunt this growth, thereby keeping individuals stuck in vulnerable size ranges. This elevates the density-independent mortality risks individuals face, ultimately threatening extinction. We saw that even once individuals are mature, they become increasingly dependent on vertebrate prey for resources for reproduction, as invertebrate prey do not provide a sufficient fraction of the adult diet to offset metabolic costs. Integrating individual-level life history processes into PVAs, as we do here, can therefore highlight potential contrasts between the effects of habitat degradation on different resources. When habitat degradation impacts both resource types, we find both destabilizing each prey group’s density-independent recruit recruitment and reducing the resource’s carrying capacity exhibit a threshold level above which extinctions risks become elevated (Fig. 6). Our results therefore suggest that in populations where individuals undergo ontogenetic niche shifts (likely the vast majority of multicellular organisms - e.g., Nakazawa 2015), habitat degradation that impacts both the primary adult and juvenile resource groups can have strong, synergistic effects on elevating extinction risks.

One potentially noteworthy result of our case study is that, in contrast to reductions in the biotic resources, habitat destruction involving the loss of nesting sites (an abiotic resource in our case study) has a small linear, rather than notably qualitatively distinct effect, in driving extinction dynamics in the spectacled caiman population at the Apaporis River. We suggest here too that PSPMs provide unique insights into this conclusion that may not have been as apparent in a PVA that ignored population size structure. For instance, the egg stage presents a notorious bottleneck in a spectacled caiman individual’s life cycle (e.g., Ayarzagüena 1983; Thorbjarnarson 1991). Indeed, this fact has been used to successfully harvest large numbers of eggs in other *Caiman* populations for decades with minimal consequences on population persistence (e.g., Brazaitis et al. 1998; Larriera and Imhof 2006; Siroski et al. 2024). Taken alone, our results are consistent with the hypothesis that the spectacled caiman population of the Apaporis River can potentially withstand even substantial reductions in the hatchling recruitment cycle. Nevertheless, our conclusions do not imply that relying on a small number of successfully nesting adult females is an adequate or even desirable conservation strategy. We highlight, for example, that our results still indicate that the destruction of nesting sites may present a non-trivial threat to population viability, especially when habitat degradation also adversely impacts recruitment in both the juvenile and adult prey groups (e.g., Figs. 6-7).

Although we applied our model to the unique/isolated population as a case study, we think our approach has broad applicability to other size-structured populations. We expect our strategy could prove quite attractive for analyzing the effects of habitat degradation on population viability for several reasons. Because PSPMs continuous measures of individual state such as somatic mass (as we have done here), they have traditionally been formulated using deterministic ordinary and partial differential equations (e.g., de Roos 1997; Diekmann et al. 2020). By contrast, PVAs rely on probabilistic conclusions. Thus, the formalisms of PSPMs may initially seem like a poor fit for PVAs. Yet we note that there is little inherent in the biological descriptions underlying PSPMs that precludes recasting them as stochastic models. Through suitable discretizations (e.g., Jagers 2010 and references herein), we aim to leverage the potential of PSPMs to link environmental perturbations to probabilistic extinction risk.

One potential advantage of our modeling approach may be essentially pragmatic. By design, PSPMs integrate the individual physiological effects of ecological processes to characterize the population- and community-level consequences of interspecific interactions. As a practical consideration, for many relatively long-lived multicellular organisms with seasonal recruitment, behavioral and physiological processes operating at the individual-level, such as metabolism and consumption in response to scarcity, etc… may often prove easier to measure than population- or community-level processes (e.g., Grimm and Railsback 2005; Sibly et al. 2013; Smallegange et al. 2017; Marques et al. 2018; Conde et al. 2019). After all, if the unit of replication is individuals, then experiments can be confined to smaller spaces, and the time-scales involved in obtaining physiological measurements may not require observing demographic turnover. Our case study involved a potential deme of a relatively well studied taxon *C. crocodilus* whose near relatives’ physiology, life-history and behavior have also been the subject of decades of close, quantitative research both in the laboratory and the field (e.g., Coulson et al. 1989; Ayarzagüena 1983; Thorbjarnarson 1991 and the references in Table 1). As a consequence, we were able to parameterize at least the autoecological components of our PSPM with reference to the literature. This might not be true of many taxa threatened by habitat degradation (e.g., Clutton-Brock and Sheldon 2010; Conde et al. 2019). Nevertheless, our approach suggests that determining measurable quantities at the individual, physiological level, and constructing models for population viability analyses from there, may, in at least some cases, prove a somewhat more feasible path than performing, for instance, multiyear field experiments to measure vulnerable populations’ demographic responses to perturbations.

We formulated our model by considering a size- and sex-structured animal population with overlapping generations. Although it may prove challenging to directly graft the model in Table (2) to conduct population viability analyses for stage-structured (e.g., holometabolous insects) or strictly parthenogenic taxa, our framework should be broadly applicable to many size-structured, multicellular species. Indeed, we suspect our basic approach can, in principle, even potentially apply to dioecious, perennial plants as well with suitable reinterpretation of the parameters (e.g., Russo et al. 2022). For instance, although somatic mass growth in plants is often limited by very different resources than animals (e.g., light), such resources are also subject to anthropogenic disturbances (e.g., Harper 1967; Crawley and Ross 1990; Lipson et al. 1999). Moreover, in the case of certain resources (e.g., pollinators, mycorrhizal fungi, etc…), the impact of habitat degradation on their population dynamics and the feedback between changing resource recruitment and population viability can be modeled quite analogously to the recruitment dynamics of prey in models (2; e.g., Holland and DeAngelis 2010) with changes in the signs of the interaction term. Thus, while we used an unique/isolated population for our case-study, we highlight modeling studies applying PSPMs to quantify extinction risks in endangered, dioecious perennial plant populations as potential future applications to evaluate the utility of our approach.

Finally, we think that one of the most substantive strengths of our approach conceptually and for conservation is its integration of feedback loops between the community consequences of habitat degradation, on the one hand, and population decline, on the other. Destabilizing prey density-independent recruitment and increasing negative prey density-dependence both reduce resource availability. Through PSPMs, we were able to characterize how reductions in prey habitat suitability cascade to drive the population viability of species at higher trophic levels by constraining somatic growth, thereby prolonging time to maturation, reducing fecundity, and keeping individuals in vulnerable size ranges for longer stretches of time and elevating starvation risk. Yet we highlight how prey availability in our model depends not only on intrinsic recruitment and habitat degradation, but also on the impact of predation from the focal species (Table 2). As the population in question declines, this can ease predation pressure on the prey resource base. More subtly, in populations undergoing ontogenetic niche shifts, changes in size-structure can further result in altered prey dynamics. Integrating feedbacks between declining prey resources and their consumer population is therefore critical to assessing habitat degradation’s adverse effects on dynamic resources in PVAs. The framework we developed and presented here uniquely enables us to account for such feedbacks between deteriorating resource availability and population vulnerability.

## Acknowledgments

The authors would like to thank E. Reysack and C. E. Wilson for helpful discussions early in the project. V. O. was supported by grants from the Undergraduate Research Opportunities Program, the College of Arts and Sciences and the Center for Applied Mathematics at the University of St. Thomas.

