## Supplementary figures and images for "Analyzing how habitat degradation drives extinction dynamics using physiologically-structured population models"

### Supplementary File S1

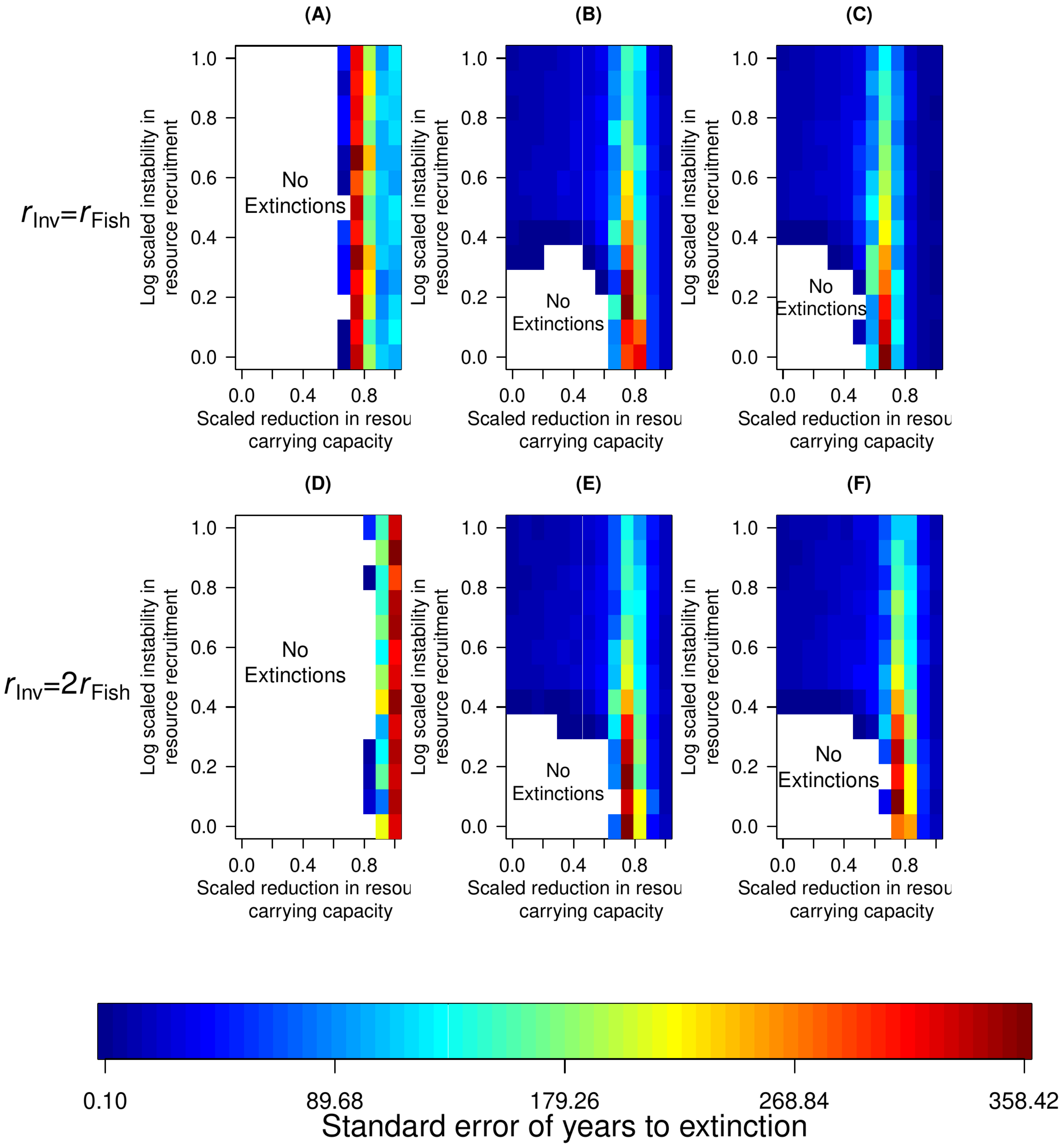

### Supplementary File S2

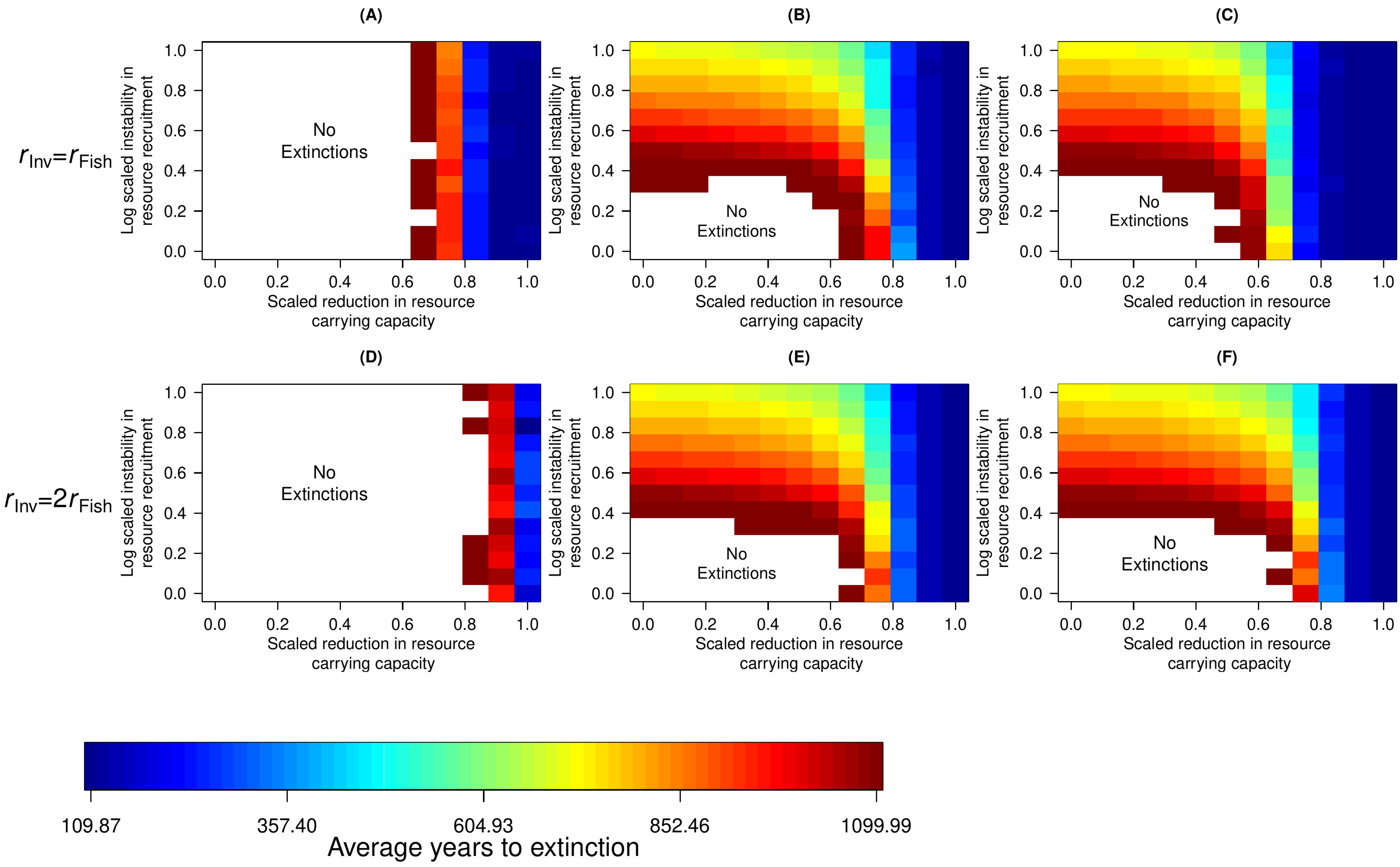
